# Population structure of the rice blast fungus *Pyricularia oryzae* in the context of elevated host heterogeneity

**DOI:** 10.64898/2026.06.16.726802

**Authors:** P-M. Marty, H. Adreit, S. Cros-Arteil, S. Guillou, S. Ravel, A. Rieux, H. Huang, L. T. Le, J-B. Morel, S. De Mita, E. Fournier

## Abstract

In a context where pathogen emergence threatens global food security, crop resistance to pathogens is crucial for preventing and controlling epidemics. Agrosystems with high intraspecific cultivated diversity are expected to better mitigate infectious diseases compared with homogeneous ones. However, the long-term impact of cultivated genotypic heterogeneity on pathogen populations is still under debate. The 1,300-year-old traditional agrosystem of the Yuanyang Terraces (YYT) in China exhibits an outstanding level of rice diversity. The rice blast fungus *Pyricularia oryzae* has been recorded in the area, making this area ideal for studying the impact of long term heterogeneous cultivation on plant-pathogen coevolution. In this study, we analysed 94 *P. oryzae* genomes from the YYT and compared them to 198 genomes representative of the worldwide diversity. We report elevated levels of genomic diversity of *P. oryzae* in the YYT. Whereas the worldwide diversity of this species is organized in four lineages, we detected seven lineages within the YYT, four of which are restricted to this area. The sampling dates of the YYT isolates (2009 to 2017) provided sufficient temporal signal to date nodes of the phylogenetic tree of isolates. Endemic lineages appeared to have arisen several centuries ago but later than the origin of the YYT themselves. We also detected recent introductions of worldwide lineages. Linkage disequilibrium analyses and *in vitro* cross experiments suggest that *P. oryzae* reproduces asexually in the YYT. These results suggest that long-term intraspecific crop diversity in the YYT has promoted the emergence and maintenance of a highly diverse, locally adapted pathogen population.

## Introduction

Modern agriculture has evolved from traditional practices through the deployment of high-yielding crop varieties over extensive areas (Zhan et al. 2015). The overall cultivated biodiversity in modern crops is lower than in traditional crops, because the genetic basis from which modern varieties are improved is limited, and because modern varieties themselves meet the criterion of genetic homogeneity (Gaut et al. 2018; Krug et al. 2023). In contrast, traditional modes of culture maintain higher levels of diversity (Hour et al. 2012). Homogeneous varieties of crops impose strong consistent selection pressures on plant pathogens, fostering the emergence and rapid spread of new virulent strains (Zhan et al. 2015; Rimbaud et al. 2018). For example, in a four-year experiment involving a resistant cultivar of oilseed rape (*Brassica napus*), the pathogenic fungus *Leptospharia maculans* overcame the resistance gene through various mutational events (Daverdin et al. 2012). Furthermore, one of the indirect consequences of plant domestication is a global decrease of plant immunity (Singh et al. 2022). In this context, pathogens impose an increasing pressure on crops, with yield losses exceeding often 20% (Savary et al. 2019).

In contrast, it has been shown that plant agrosystems that are diversified interspecifically (agroforestry, cover crops, intercropping, rotation) or intraspecifically (cultivar mixture) provide numerous ecosystem services (more stable yields, soil quality, protection against diseases and pests, general resilience, see Beillouin et al. 2021; Mathieu et al. 2025, Vialatte et al. 2025). In particular, genetic and spatial heterogeneity of crops is a source of resistance durability (Karasov et al. 2020), primarily by imposing a more complex adaptive landscape to pathogens and thereby restricting their evolutionary ability. Other factors come into play such as dilution of compatible host, resistant hosts acting as barrier, and plant-plant interactions increasing immune response of individual plants (Zhan et al. 2015; Borg et al. 2018; Mathieu et al. 2025).

Here we are primarily interested in the evolution of pathogens in a diversified host environment. Such evolutionary mechanisms of resistance durability are more difficult to tackle than in homogeneous environments as it requires considering longer time frame and investigating the population structure of pathogens in parallel to host diversity in a given agrosystem.

We focused on an agricultural rice-growing system with high levels of intraspecific crop diversity maintained over centuries. The Yuanyang Terraces (YYT), located in the Yunnan province in Southern China, are a traditional agrosystem where rice has been cultivated in irrigated paddies since the 9th century (Jiao et al. 2012, Hua & Zhou 2015). Traditional agricultural practices involve about 200 traditional varieties (Jiao et al. 2012), and the two subspecies of Asian cultivated rice subspecies (*Oryza sativa* subsp. *indica* and *O. sativa* subsp. *japonica* respectively) are grown in this system (Liao et al. 2016). A non-market seed exchange system explains the in-situ maintenance of a very high level of cultivated biodiversity (Hannachi & Dedeurwaerdere 2018). Traditional varieties, or landraces, cultivated in YYT are population varieties exhibiting more diversity than modern varieties (He et al. 2011, De Mita et al. 2024, Zhu et al. 2025), as exemplified by the emblematic indica landrace Acuce that exhibits 38% of the diversity of the entire indica subspecies (De Mita et al. 2024). This diversity is at least partially functional with respect to pathogen resistance since immune receptors exhibit more extensive gene repertoires and increased nucleotide diversity with traces of balancing selection in the YYT compared to modern varieties (Gladieux et al. 2024). From the epidemiological point of view, no severe epidemics have been reported in the YYT, where rice diseases cause less than 1% yield loss (Sheng 1990), as compared to around 30% at the worldwide scale (Savary et al. 2019).

Among several fungal diseases recorded in the YYT, *Pyricularia oryzae* (syn. *Magnaporthe oryzae*) is responsible for the blast disease on *Poaceae* (Gladieux et al. 2018a). Previous studies of the diversity of *P. oryzae* on rice identified four lineages at the worldwide scale (WW1 to WW4), and showed that South-East Asia was the centre of origin of the disease on rice (Saleh et al. 2014; Gladieux et al. 2018b; Latorre et al. 2020; Thierry et al. 2022, reviewed in Baudin et al. 2024). Two studies focusing on population structure of *P. oryzae* in the YYT have revealed a surprising diversity for this fungus compared to worldwide diversity. Using microsatellites, it was shown that *P. oryzae* populations segregated in two groups locally adapted to indica and japonica landraces, respectively (Liao et al. 2016). A second study based on whole genome data (Ali et al. 2023) showed that the *P. oryzae* genetic group associated to indica landraces was itself subdivided into five lineages, three of which being endemic to the YYT. However, we still lack an integrated view of how blast genetic diversity is organized in this peculiar agrosystem.

Based on phylogenomic analyses assuming a molecular clock, it has been suggested that worldwide lineages (WW1-4) emerged in the 8th century (Gladieux et al. 2018b), approximately at the same period than the establishment of the YYT agrosystem. The age of YYT-specific lineages is currently unknown. They may have diverged from external populations introduced at the moment when the YYT agrosystem was established, possibly through co-dispersion with their rice hosts. If so, they would have existed throughout the whole history of the YYT, possibly with later diversification. Alternatively, the extant YYT lineages may be the result of more recent colonization events of the area. Understanding how the *P. oryzae* population evolves in this system would help predicting the effect of host diversity on the long-term dynamics of pathogen populations. Testing this hypothesis requires extensive sampling of *P. oryzae* in the YYT to replicate phylogenetic analysis allowing dating of divergence nodes.

Besides, previous studies showed that the reproduction regime of *P. oryzae* on rice differs across the known worldwide genetic lineages, with WW1 exhibiting signatures of sexual reproduction and WW2-4 being strictly clonal (Saleh et al. 2012, 2014; Gladieux et al. 2018b; Latorre et al. 2020; Thierry et al. 2022). The reproductive regime impacts the emergence of new allelic reassortments among genotypes, hence the response to selection. Understanding how *P. oryzae* populations reproduce in the YYT is therefore crucial.

In this study, we aimed to better understand how the elevated level of cultivated biodiversity of rice in the YYT impacts the population structure and evolution of the blast fungus. We sampled additional isolates within the YYT and combined population genomics with *in vitro* crossing experiments to address the following questions: 1) how many genetic lineages are recorded in the YYT and how are they related to each other? 2) what is the tempo and timing of emergence of these lineages? 3) what is the mode of reproduction of *P. oryzae* in the YYT?

## Materials and methods

### Datasets, sampling, DNA extraction and genome sequencing

In the present study, we used 292 *P. oryzae* isolates from rice, of which 94 were sampled in the YYT and 198 were sampled from worldwide (mostly Asian) populations outside of YYT. Fungal isolates were recovered from samples collected on blast symptomatic rice plants mostly belonging to cultivated Asian rice subspecies (*Oryza sativa*), with four exceptions sampled on *Oryza glaberrima* and *O. longistaminata* (Table S1). Out of the 292 isolates, 203 have already been fully sequenced and used in former studies (Gladieux et al. 2018b; Pordel et al. 2021; Were et al. 2021; Thierry et al. 2022; Ali et al. 2023; Le Naour-Vernet et al. 2023; Lassagne et al. 2024). For the 89 new isolates, single spores of *P. oryzae* were isolated from colonies placed in humid chamber at 21°C for 1–2 days. Fungal isolates were then grown on rice flour medium following Nottéghem & Silué (1992), and stored on filter paper at −20°C following Valent et al. (1986). DNA extractions were performed following Ali et al. (2023). The 89 new genomes generated for this study were acquired using Illumina HiSeq 2500 or Illumina NovaSeq 6000, with paired-end reads of length 150 bp. Two *P. oryzae* isolates from *Stenotaphrum* were used as outgroup (Table S1). Complete information about the geographic origin, date of collection, sequencing technology, available sequencing datasets and references are provided in Table S1.

### SNP calling and filtering

Mapping of all reads from the 292 samples was performed separately for each isolate against the version 8 of the *P. oryzae* 70-15 genome reference (Dean et al. 2005), masked for the mitochondrial genome. The quality control, mapping and post-processing alignment were performed using RattleSNP (https://rattlesnp.readthedocs.io/). RattleSNP uses FastQC v. 0.12.1 to assess the quality of the FastQ, BWA MEM v. 0.7.17 to perform the mapping, and SAMtools v. 1.21 (Danecek et al. 2021) to index BAM files, flag reads (e.g. reads duplicates) and calculate sequencing depth and coverage.

Sequencing depth varied among isolates from different source. As sequencing depth greatly impacts mapping depth, SNP calling and filtering of VCF were performed with a custom pipeline aiming to take into account the variation of mapping depth among isolates (https://gitlab.cirad.fr/phim/pmarty/rattlepm). In this pipeline, SAMtools was used to remove reads duplicates from BAM files (markdup-r option). SNP calling was performed using BCFtools v. 1.16 (Danecek et al. 2021) with the following parameters: maximum mapping depth of 500 reads per position, Phred-Score > 30 and indels skipped. In the next step we excluded sites with mapping depth deviating from the genome-wide distribution. Mapping depth per site was calculated using the site-mean-depth option of VCFtools v. 0.1.16. For each isolate, we identified outliers sites using the boxplot.stat() function of R 4.4.1, computed mean (µ) and standard-deviation (σ) of mapping depth without outliers, and kept only sites with a mapping depth comprised in [µ +/- 3σ] using the – minDP and –maxDP options of VCFtools.

Filtered individual VCF were then merged into a single VCF with merge-all option of BCFtools. This merged VCF was then filtered using VCFtools with criteria: site quality (Q) ≥ 29, missing data=1 (no missing data allowed), and no minor allele frequency filter. Indels and sites with more than two alleles were filtered out.

### Genealogical relationships

VCF with only biallelic sites were converted into FASTA using vcf2phylip (https://github.com/edgardomortiz/vcf2phylip). Two alternative tools making different assumptions were employed. Genealogical networks were inferred with SplitsTree v. 4.19.2 (Huson & Bryant 2006) using default parameters. Such networks accommodate potential recombination in the forms of reticulation of the network. In parallel, we reconstructed total evidence maximum likelihood genealogical trees with RAxML-NG v. 1.1.0 (Kozlov et al. 2019) using the GTR+Γ model with 100 replicates for bootstrapping. Phylogenetic methods such as RAxML assume that no recombination occurred in the history of the sample. Two *P. oryzae* isolates from *Stenotaphrum* sp. were used as outgroup to root the genealogical tree.

### Population subdivision

We performed three different analyses to infer population subdivision. Firstly, we conducted an unsupervised analysis of the repartition of the global genetic variance into several groups by running a Principal Components Analysis (PCA) using the R package Adegenet (Jombart 2008). Secondly, we use the unsupervised clustering algorithm sNMF (sparse non-Negative Matrix Factorization) implemented in the R package LEA (Frichot et al. 2014). We let K (the number of inferred genetic clusters) vary from 2 to 12, ran 50 iterations with 10 repetitions each for each value of K, and selected the best repetition based on the lowest cross-entropy. This hypothesis-based algorithm determines the composition of the K groups by minimizing the linkage disequilibrium within each group, and provides estimates of the coefficients of ancestry of each individual in each of the K groups for each value of K. Thirdly, we conducted a DAPC (Discriminant Analysis of Principal Components) using the R package Adegenet (Jombart et al. 2010). We let K (the inferred number of genetic groups) vary from 2 to 12, performed 10 repetitions, and selected the best repetition (identified by the lowest Bayesian Information Criterion). This semi-supervised, hypothesis-free algorithm first reduces the number of dimensions using a PCA, then uses a DA (Discriminant Analysis) to maximize the inter-group genetic variance and minimize the intra-group genetic variance. We retained the principal components and discriminant functions that explained most of the variance. Isolates were assigned to a given cluster when the corresponding assignation probability was at least 50%. We examined the consistency of inferred clusters with previously structure analyses from the literature, for the isolates with the available information. Other isolates were then assigned to previously described lineages or new lineages if necessary.

### Molecular dating

Since samples were collected a different dates, it is theoretically possible to assume a molecular clock and estimate the age of ancestral nodes based on temporal signal present in the data (Rieux & Balloux 2016). Considering that the worldwide sample may be affected by large-scale history potentially blurring the temporal signal, we restricted this analysis to the YYT isolates, accounting for 94 samples.

#### Filter of recombining regions

To reduce the effect of recombination in phylogenetic analysis, we used Gubbins (Croucher et al. 2015) to predict the recombining regions within each of the seven chromosomes of *P. oryzae* (based on the reference genome 70-15). Gubbins detects recombinant regions as genomic segments enriched in clustered substitutions whose phylogenetic signal is incompatible with the whole-genome genealogy. We used the genealogy inferred from the 94 YYT isolates as starting tree. We configured Gubbins to use the RAxML-NG software to build iterative trees. We set minimum and maximum window size to 0.2 and 20 kb, respectively, and used the default parameter of maximum 5 iterations. Regions identified as recombinant by Gubbins were filtered using a custom script.

#### Detection of temporal signal

We used Phylostems (Doizy et al. 2023) to identify clades exhibiting temporal signal. Phylostems investigates temporal signal within every clade of a tree by fitting a linear regression of the number of substitutions accumulated from the root to the tips of a clade as a function of sampling dates. We used the genealogy inferred from the 94 YYT isolates from non-recombinant regions rooted with the two outgroups, and retained the clades for which the linear regression was significant and grouping a maximum of isolates.

#### Estimation of mutation rate with BEAST

We used BEAST v1.8.4 (Drummond & Rambaut 2007) to estimate the mutation rate within subtrees exhibiting significant temporal signal as detected by Phylostems. We first generated the XML input file using BEAUti with the following options: include tip dates, GTR + Γ + invariants sites model, uncorrelated relaxed clock (with lognormal relaxed distribution), Extended Bayesian Skyline Plot, and a uniform prior on mutation rate bounded by 10^-9^ and 10^-5^. This range includes mutation rates estimated for several Ascomycota, including *P. oryzae* (Kasuga et al. 2002; Demené et al. 2016; Gladieux et al. 2018b; Latorre et al. 2020). Following Rieux & Balloux (2016), in order to save computing time, we used the VCF file containing only SNPs and specified the numbers of invariant sites A, T, C and G in the XML input file. Next, BEAST v1.8.4 (Drummond & Rambaut 2007) was used to estimate the mutation rate for each subtree. To check convergence, we ran five independent chains, in which samples were drawn every 10,000 MCMC steps from a total of 100,000,000 steps, after a discarded burn-in of 1,000,000 steps following Rieux & Balloux (2016). After checking for convergence, we randomly selected one chain for each subtree and merged posterior distributions of the mutation rate parameter from these chains, thereby generating a composite posterior distribution that captures the variability between subtrees. Estimates of the mean (µ) and variance (σ) of the estimated mutation rate were obtained by fitting a normal law to this composite distribution.

#### Datation of genealogical nodes with BEAST

The estimated mutation rate previously obtained was used to estimate node ages in the genealogy of the 94 YYT isolates. BEAUti was used to generate the XML input file with the following options: include tip dates, GTR + Γ + invariants sites model, uncorrelated relaxed clock (with lognormal relaxed distribution with parameters µ and σ previously estimated), Extended Bayesian Skyline Plot. Specifying invariant sites and running BEAST analysis were performed as in the previous section. The time-scaled tree was visualized using the R package ggtree (Yu et al. 2017).

### Migration inference

To describe the ancestral relationships between lineages and the signature of potential gene flow events among lineages, we studied the patterns of population divergence using TreeMix (Pickrell & Pritchard 2012), including 267 isolates that were assigned to a lineage with more than 50% ancestry proportion (sNMF analysis) and the two *Stenotaphrum* isolates as outgroup. We ran TreeMix by letting the number of inferred migration events (*m*) vary from 1 to 10. We performed 15 repetitions for each value of *m*, with the size of the block of SNP considered in one repetition (*k*) ranging from 50 to 120 with steps of 5. The optimal number of migration events was inferred using OptM (Fitak 2021).

### Diversity and divergence

Statistics of genetic diversity and divergence were calculated within and between YYT lineages using EggLib 3.3.4 (Siol et al. 2022) on the YYT dataset with biallelic and invariant sites. We estimated the level of polymorphism within each lineage with two statistics: per site nucleotide diversity (*π*) and Watterson’s estimator of *θ* (*θ_w_*). We computed Tajima’s *D*, a statistic that measures the departure from neutrality. Population differentiation was estimated using Weir and Cockerham’s *F_ST_*.

### Linkage disequilibrium (LD) decay and inference of recombination

For each lineage including more than 3 samples, the pairwise linkage disequilibrium (*r²*) was calculated for all pairs of SNPs using EggLib. Since recombination leads to a decrease of LD with increasing physical distance between markers, we represented binned *r²* as a function of distance between pairs of SNPs. We grouped *r²* for classes of distances of size increasing with the distance to compensate the decreasing number of available pairs of sites, with a size varying from 300 bp to 20,000 bp. We represented the mean *r²* within classes against the class midpoints.

We also performed the PHI test (Pairwise Homoplasy Index) implemented in SplitsTree to test for the null hypothesis of strict clonality. This test assesses pairwise homoplasy, and the null hypothesis of no recombination is tested using random permutations of the positions of the SNPs based on the expectation that sites are exchangeable if there is no recombination (Bruen et al. 2006).

### *In silico* mating-type detection

For each YYT isolate, raw reads were mapped separately against the sequences of MAT1.1 and MAT1.2 mating type loci (Kanamori et al. 2007) using the RattleSNP workflow, setting the-a option for depth in SAMtools in order to include positions with zero coverage. Sites were classified in three categories according to mapping depth: depth < 10 (low), 10 < depth < 300 (acceptable), depth > 300 (high). Sites with high depth may be caused by reads erroneously mapped due to sequence similarity, therefore we counted only reads falling in the two other categories. Isolates were assigned to the reference MAT sequence for which most reads fell into the acceptable mapping depth category while no reads fell into this category for the alternative MAT sequence, consistent with the haploid, heterothallic nature of the fungus.

### *In vitro* sexual crosses experiments

In order to test whether YYT isolates were capable of sexual reproduction, we conducted *in vitro* sexual crosses between all compatible pairs of YYT isolates (i.e. between isolates carrying opposite mating types), as well as *in vitro* sexual crosses between each YYT isolate and a fertile reference isolate with opposite mating type. The fertile isolates used as reference were CH0999, carrying mating type MAT1.1, and CH0997, carrying mating type MAT1.2 (Saleh et al. 2012; Lassagne et al. 2022). Each isolate was first grown on rice flour medium (Nottéghem & Silué 1992) for 5-6 days until the Petri dish was well-covered with sporulating mycelium. For each crossing test, one plug of mycelium (5 mm²) was sampled for each selected isolate and transplanted on the a new Petri dish with rice flour medium. The culture was incubated at 21°C for three weeks. All combinations of crosses were performed by setting mycelia plugs from 4 isolates on the same Petri dish as in Saleh et al. (2012). Presence/absence of perithecia at the confrontation lines between mycelia of adjacent isolates was scored under stereo-microscopy after three weeks of culture.

## Results

### Genome sequencing and SNP calling

Reads from 203 isolates from the literature and 89 newly-sequenced isolates were included in the analysis. The mean sequencing depth per isolate ranged from 6 to 319X (Table S2). After mapping and filtering for over-mapped sites, the mean mapping depth ranged from 6X to 253X per isolate (Table S3). SNP calling led to the identification of 41,024 biallelic SNPs for the dataset of 292 worldwide isolates, and of 20,234 biallelic SNPs for the dataset of 94 YYT isolates.

### Lineage assignation in the YYT: three worldwide lineages and four endemic lineages

We first inferred the population structure of the full dataset (n=292) using sNMF. The selection of the number of clusters was not straightforward, since the cross-entropy signal decreased monotonically up to K=20, but the rate of decrease showed an inflexion at K=5 (Figure S1). Importantly, K=5 was the first value at which previously defined worldwide lineages were nearly perfectly recovered (Figure 1A, Figure S1). The 133 isolates previously assigned to worldwide lineages WW1-4, were mostly assigned to four matching clusters, with 6 exceptions. The exceptions were 5 isolates not assigned at the 50% threshold and a single isolate assigned to an inconsistent cluster (CH0452, assigned to the fifth cluster along YYT isolates; Table S4). According to the correspondence between sNMF clusters and worldwide lineages, we attributed 67 previously unassigned lineages to one of the four worldwide clusters (Table S4). In contrast, isolates previously assigned to YYT1-3 lineages were not separated accordingly in our sNMF analysis (Figure 1A), even with higher K values (Figure S1).

**Figure 1.**
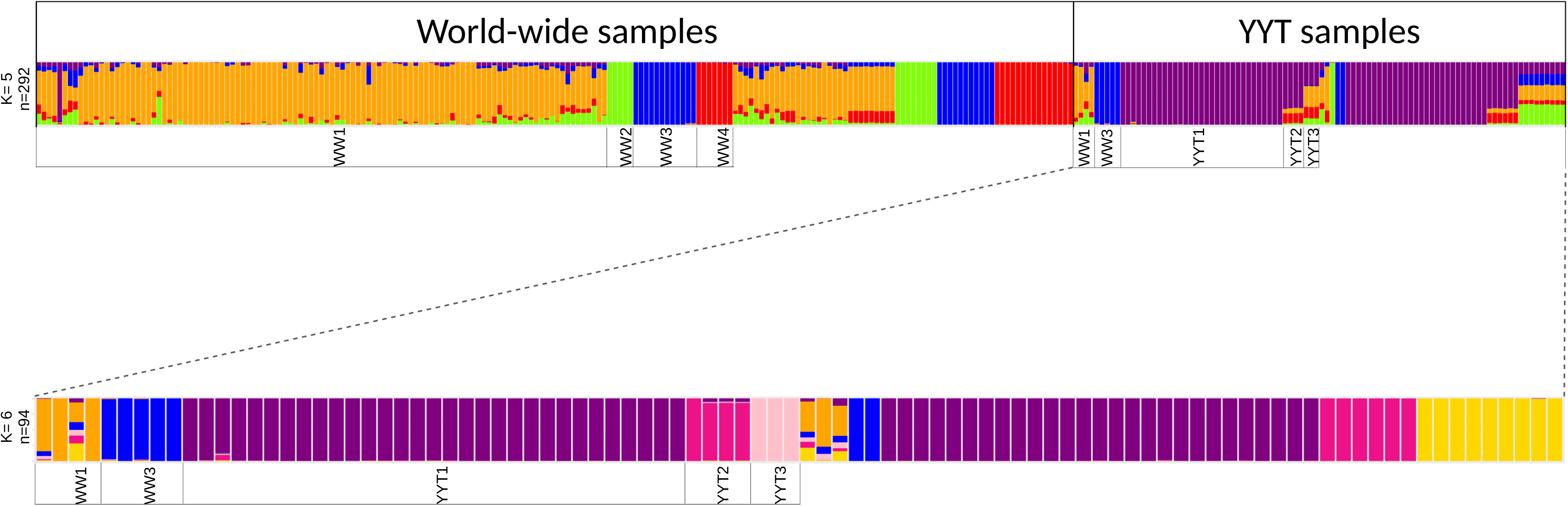
Population structure inferred with sNMF for the whole dataset (n=292 isolates, 41,024 SNPs) and for the dataset restricted to YYT isolates (n=94 isolates, 20,234 SNPs). The top panel shows the population structure at K=5 for the whole dataset, the bottom panel shows the population structure at K=6 for the YYT dataset. In both panels, each isolate is represented by a vertical bar plot subdivided into K segments corresponding to the probability of ancestry of the isolate in each of the K groups. Assignments of isolates established from previous studies (Gladieux et al. 2018b; Latorre et al. 2020; Thierry et al. 2022) are labelled.

Considering that the imbalanced representation of the different lineages in the full dataset (with the domination of the WW1 lineage outside of the YYT and of the YYT1 lineage within the YYT) could have obscured signal segregating YYT lineages, we ran sNMF for the dataset restricted to the 94 YYT isolates. The cross-entropy signal reached a minimum between K=5 and K=7 (Figure S2). At K=6, isolates previously assigned to lineages YYT1-YYT3 were consistently recovered as three distinct sNMF clusters (Figure 1B). Based on this result, we assigned 33 previously unassigned isolates to lineages YYT1 and YYT2, but none to YYT3 (Table S4). Two of the other sNMF clusters corresponded to worldwide lineages WW1 and WW3, consistently to the analysis with the full dataset (which the exception of CH2307, the unique representative of WW2 which was left unassigned). Finally, a group of 9 isolates, all sampled on japonica rice and unassigned up to this point, were assigned to the last cluster, which we therefore interpreted as a previously unknown lineage and named YYTJ. Overall, seven clusters were recovered in the YYT (Table S4): four endemic (YYT1: 58 isolates, YYT2: 10 isolates, YYT3: 3 isolates, YYTJ: 9 isolates) and three worldwide lineages (WW1: 6 isolates, WW2: 1 isolate, WW3: 7 isolates). We replicated the full structure analysis using DAPC and obtained consistent results, although DAPC reported less conservative assignment probabilities (Figures S1 and S2, Table S4). Following DAPC results, we assigned all remaining isolates to WW1 (Table S4).

### Relationships between lineages

The seven lineages identified in our dataset were clearly delineated on the genealogical network built from the complete dataset (Figure 2A). All lineages except WW1 were connected to the rest of the network by an edge with minor reticulations, suggesting monophyletic ancestry. WW2, WW3, WW4 and YYTJ appeared to stem from the heavily reticulated central part of the network while YYT1 and YYT2 were grouped together, with YYT3 in an intermediate position (grouped with the YYT1-YYT2 group by a mildly reticulated branch). The closest worldwide lineage for the YYT1-YYT3 group was WW4, which was interestingly absent from the YYT isolates, with a large reticulation at their common base, suggesting that they had a common ancestor who experienced recombination. Topology of the RAxML-NG genealogy was congruent with the genealogical network (Figure S3).

**Figure 2.**
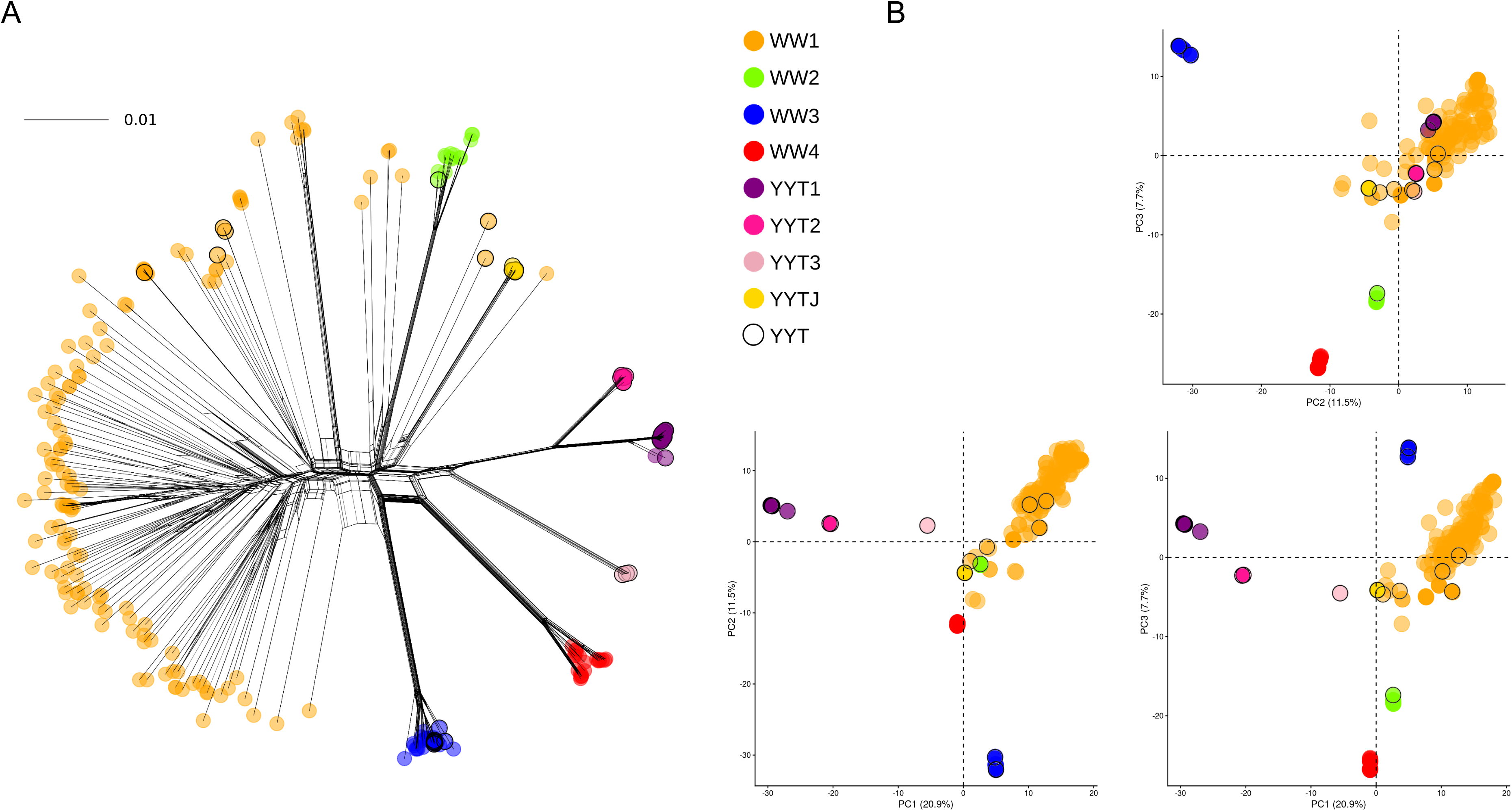
Population structure inferred with genealogical network and PCA for the whole dataset (n=292, 41,024 SNPs). A. Genealogical network inferred with SplitsTree. Scale bar represents the number of substitutions per site. B. Principal component analysis (PCA) with the first three PC axes. Isolates are coloured according to their lineage attribution deduced from sNMF analyses. Black circles pinpoint YYT isolates.

The PCA (Figure 2B) confirmed that the seven lineages were separated from each other and identified a major component (PC1: 20.9% of the total variance) separating lineages YYT1 and, successively, YYT2 and YYT3 on one side and the bulk of WW1 isolates on the other, with YYTJ, WW2-WW4 and part of WW1 being intermediate. PC2 and PC3 seemed to capture differentiation of worldwide lineages, with all YYT endemic lineages being closed to the origin, mixed with WW1 isolates.

The same analysis genealogical network reconstruction and PCA analysis was performed on the YYT dataset (n=94) alone, supporting the same conclusions when applicable (Figure S4). Lineages YYT1, YYT2 and YYT3 grouped together in the genealogical network, with YYT3 appearing as intermediate with mild reticulations. These three lineages were separated from the rest of samples on the first PC of the PCA (18.7% of the total variance). YYTJ formed a single cluster among the few WW1 isolates. WW3 was isolated from the rest both on the network and on PC2 (11.4% of the total variance). The only new observation was that PC3 (10.3% of the total variance) was explained by the divergence between YYTJ and WW1 isolates, which can be explained by the fact that WW4 was absent and WW2 was represented by a single isolate. Genetic network and genealogical tree inferred with RAxML-NG showed concordant topologies (Figure S5). Therefore, the previously known lineages YYT1, YYT2 and YYT3 seem to have a common origin with respect to the WW1 lineage which locates at the centre of the network. By contrast, the new lineage YYTJ seem to have emerged independently from WW1.

### A single migration event between WW1 and YYT3

We searched for signatures of hybridization events between lineages, using TreeMix (Pickrell and Pritchard 2012). We analysed 267 isolates that could be assigned to a lineage with two *Stenotaphrum* isolates as outgroup and treated worldwide and endemic lineages as populations. The optimal number of migrations events was *m*=1, suggesting that only one hybridization event occurred from WW1 and YYT3 (Figure S6). This migration event was consistently predicted in each of the 15 repetitions for *m*=1.

### Endemic lineages are estimated to have diverged from worldwide lineages several centuries after YYT construction

After filtering recombinant regions identified with Gubbins for YYT isolates, we obtained a dataset comprising 24,830,644 invariant sites and 17,149 SNPs. Merging this YYT dataset with the two *Stenotaphrum* isolates as outgroups resulted in a dataset with 58,909 biallelic SNPs without missing data. The genealogy reconstructed using this last dataset was identical to the genealogy obtained with the whole set of SNPs. Taking into account the date of sampling, we identified three subtrees exhibiting significant temporal signal, and we independently performed tip-dating estimation of the mutation rate for each subtree (Figure S7). The combination of the three marginal posterior distributions of the substitution rates fitted a normal distribution with a mean of 2.07×10⁻⁷ and a standard deviation of 1.24×10⁻⁷ substitutions/site/year. Although the use of this composite distribution may decrease the precision of mutation rate estimates, it offers a conservative approach for subsequent analyses by integrating variability across different clades of the global tree.

This composite distribution of the substitution rate parameter was used as prior to perform rate-dating on the whole set of YYT isolates. The YYTJ endemic lineage diverged from WW1 around year 1461 AD with a large credibility interval (95% highest posterior density [HPD]: 650–1900 AD). The three other endemic lineages formed a monophyletic group that diverged around year 1188 AD, with a very large credibility interval (95% HPD: 200–1900 AD). Lineage YYT3 segregated from this common ancestral population in year 1493 AD (95% HPD: 780–1900 AD), YYT2 in year 1923 AD (95% HPD: 1795–1995 AD), and YYT1 in year 1943 AD (95% HPD: 1850–2000 AD). The four YYT endemic lineages had therefore a multi-century history, but they likely diverged from worldwide lineages after the origin of the YYT agrosystem.

### Blast representatives of worldwide lineages arrived in the YYT within the last two centuries

Assignment of YYT isolates to WW2 and WW3 was unambiguous (Figure 1). Furthermore, for each of these clonal lineages, YYT isolates perfectly grouped with worldwide isolates. Previous population genomics analyses (Gladieux et al. 2018b; Lattorre et al. 2020) dated the clonal expansion of these two lineages within the last 200 years. Therefore, YYT isolates belonging to these lineages were likely introduced recently (i.e. within the last two centuries) in the YYT through migration. YYT isolates assigned to WW1 are polyphyletic in the genealogy (Figure 2, Figure 3, Figure S3), which could be due to multiple introductions.

**Figure 3.**
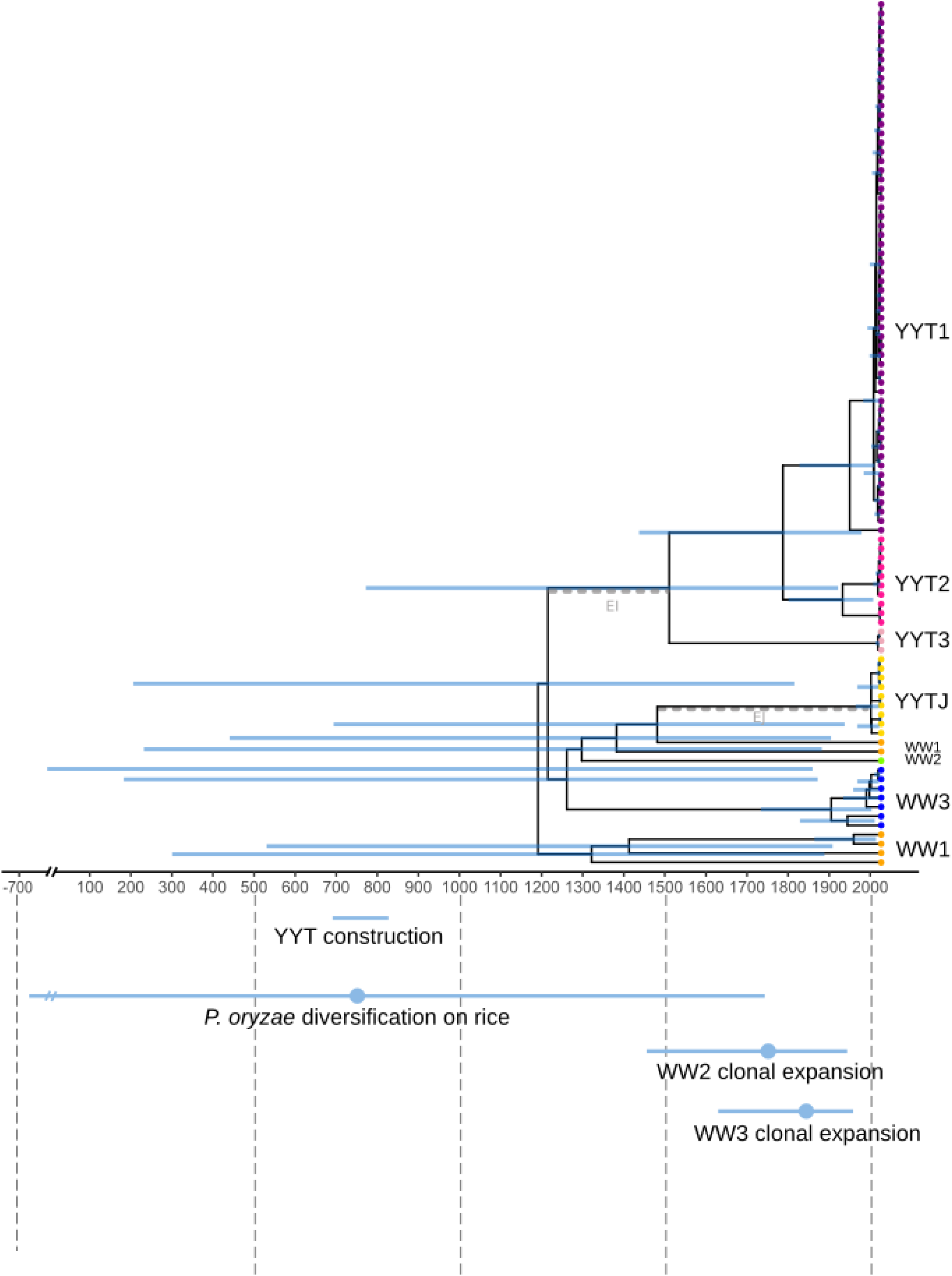
Tip-calibrated genealogy inferred by maximum-likelihood phylogenetic inference in the YYT dataset (n=94 isolates, 20,234 SNPs). Blue bars show the 95% highest probability density of nodes ages. Colour of tips represents isolate assignation to lineages deduced from sNMF analyses. Grey dashed lines pinpoint branches with emergence events leading to YYT endemic lineages on indica rice (EI) and on japonica rice (EJ), respectively. Approximate dates or historical periods related to the history of the YYT or to the evolutionary history of *P. oryzae* on rice at the worldwide scale are indicated for context.

Altogether, our results suggest a complex, multi-steps demographic history of *P. oryzae* lineages in the YYT, with two ancient events of local emergence of endemic lineages (one of which followed by the diversification of three endemic lineages on indica landraces), and probable recent introductions of representative of clonal lineages.

### Diversity within and between lineages in the YYT

The 7 lineages present in the YYT exhibited substantial variation in genetic diversity (Table 1). Nucleotide diversity was highest in WW1 (*π*=1.86×10^-4^, *θ*_W_=2.01×10^-4^), and approximately one order of magnitude lower in WW3 (*π*=1.93×10^-5^*, θ*_W_= 2.04×10^-5^). Endemic YYT lineages displayed consistently low levels of diversity (*π* ranging from 9.99×10^-6^ for YYT2 down to 7.5×10^-7^ for YYT3*, θ*_W_ ranging from 7.99×10^-6^ for YYT2 down to 7.50×10^-6^ for YYT3). However, the small number of isolates for YYT3 (n=3) limits the reliability of diversity estimates for this lineage. Tajima’s *D*, which contrasts *π* with *θ*_W_, revealed marked differences in allelic frequency spectra among lineages, suggesting contrasted demographic or selective regimes. The worldwide lineages WW1 (n=6) and WW2 (n=7), as well as the endemic lineages YYTJ (n=9) and YYT3 (n=3), displayed Tajima’s *D* values close to zero or slightly negative, patterns that do not depart strongly from neutral expectations and may reflect demographic stability or weak purifying selection. In contrast, YYT1 (n=58) exhibited a strongly negative Tajima’s *D* (*D*=-2.2) consistent with a recent population expansion or a directional selection regime, whereas YYT2 (n=10) exhibited a strongly positive Tajima’s *D* (*D*=1.3) consistent with a recent population contraction or balancing selection. This pattern is indicative of contrasted demographic and selective dynamics among lineages.

**Table 1.** Genetic diversity within the 6 lineages recorded in the YYT with at least 3 isolates. *N*: sample size, *L*: total number of sites analyzed, *S*: number of segregating sites, *π*: per-site nucleotide diversity, *θ*_W_: per-site Waterson’s estimator of *θ*, *D*: Tajima’s D.

| Lineage | N | L | S | $\pi$ | $\theta_w$ | *D* |
| --- | --- | --- | --- | --- | --- | --- |
| YYTJ | 9 | 25,785,090 | 345 | $4.733 \times 10^{-6}$ | $4.923 \times 10^{-6}$ | -0.2004 |
| YYT1 | 58 | 25,785,090 | 1017 | $3.188 \times 10^{-6}$ | $8.520 \times 10^{-6}$ | -2.2498 |
| YYT2 | 10 | 25,785,090 | 583 | $9.988 \times 10^{-6}$ | $7.992 \times 10^{-6}$ | 1.2514 |
| YYT3 | 3 | 25,785,090 | 29 | $7.498 \times 10^{-7}$ | $7.498 \times 10^{-7}$ | 0 |
| WW1 | 6 | 25,785,090 | 12725 | $1.860 \times 10^{-4}$ | $2.010 \times 10^{-4}$ | -0.4535 |
| WW3 | 7 | 25,785,090 | 1204 | $1.935 \times 10^{-5}$ | $2.045 \times 10^{-5}$ | -0.3505 |

Weir and Cockerham’s *F_ST_* computed for all pairs of YYT lineages (comprising at least three isolates) were always above 0.5, and above 0.8 for pairs not involving WW1 (Table S5), showing that divergence was maintained among lineages despite the fact that they occur in sympatry. The smaller values for pairs involving WW1 can be explained by the fact that WW1 is significantly more diverse than the other lineages.

### Loss of sexual reproduction in YYT

The *in silico* determination of mating types for the 94 YYT isolates was unambiguous, with a single mating type per isolate (as expected since *P. oryzae* is heterothallic). Among the six YYT isolates assigned to WW1, both mating types were represented (ratio MAT1.1:MAT1.2=4:2). In contrast, the other lineages were fixed for a single mating type (MAT1.1 for lineage YYTJ, WW2 and YYT3; MAT1.2 for YYT1, YYT2 and WW3) (Table 2). The global ratio MAT1.1:MAT1.2 (17:77) was unbalanced and unexpected to be observed by chance assuming balanced frequencies (χ² =38.3, P-value=6×10^-10^).

**Table 2.** Distribution of mating types within each of the 7 lineages recorded in the YYT.

| Lineage | N | Mat1.1 | Mat1.2 |
| --- | --- | --- | --- |
| YYTJ | 9 | 9 | 0 |
| YYT1 | 58 | 0 | 58 |
| YYT2 | 10 | 0 | 10 |
| YYT3 | 3 | 3 | 0 |
| WW1 | 6 | 4 | 2 |
| WW2 | 1 | 1 | 0 |
| WW3 | 7 | 0 | 7 |

To further investigate footprints of recombination within each lineage comprising at least three isolates, we computed the LD decay for each lineage present in the YYT with more than 3 individuals. A decay of LD was detected for WW1, YYT1 and YYTJ but not for WW3 and YYT2 (Figure 4). The decay signature was continuous for WW1 and mostly caused by a decrease from the first to the second window for YYT1 and YYTJ. PHI tests (performed within each lineage with at least 3 isolates) significantly rejected the null hypothesis of clonality for WW1 and YYT2, and suggested clonality for the other lineages tested (WW3, YYT1, and YYTJ; Table S6). The level of evidence was much more limited for YYT2 than WW1 (P-value=0.0428). Taken together, these results pointed to unambiguous recombination signatures in the worldwide lineage WW1 only.

**Figure 4.**
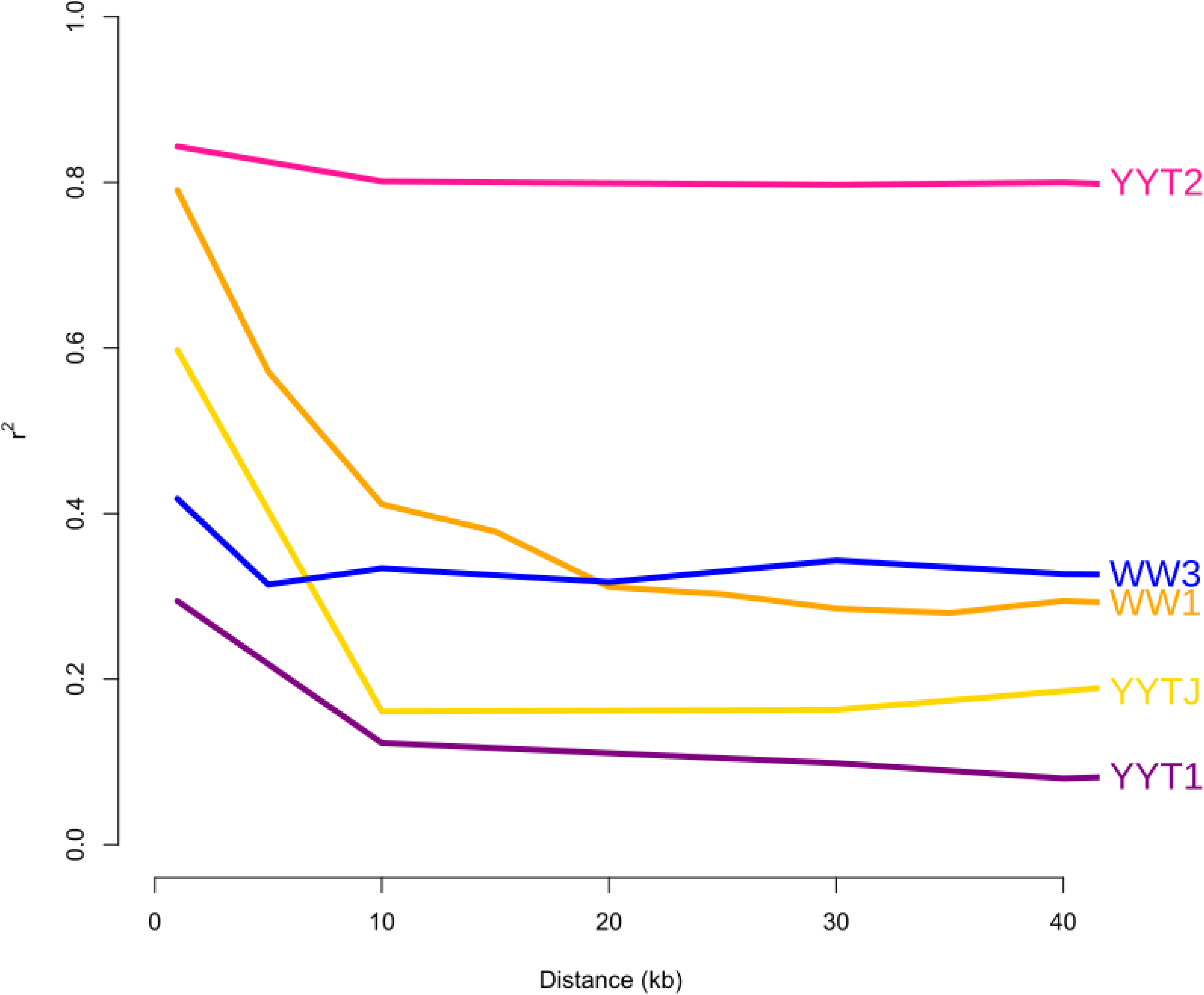
Linkage disequilibrium as a function of physical distance between markers within lineages with more than three isolates. Colours correspond to lineages delineation deduced from sNMF analyses.

To test the ability for sexual reproduction of *P. oryzae* in the YYT, we performed *in vitro* crosses between all compatible pairs of YYT isolates (i.e. carrying opposite mating types) on one hand, and between each YYT isolate and a fertile (non-YYT) reference isolate of opposite mating type as a control on the other hand. We observed no perithecia production among the 1309 YYT x YYT crosses. Out of the 94 YYT x fertile reference crosses, 85 produced perithecia, but only on the reference side, indicating that YYT isolates were male-fertile (able to induce perithecia production in an isolate of opposite mating-type) but female-sterile (unable to produce perithecia). However, seven YYT isolates (belonging to five different lineages: 1 isolate from YYTJ, 1 from YYT2, 3 from YYT3, 1 from WW1 and 1 from WW3) did not induce perithecia production in the reference isolate, suggesting that they are male-sterile, and two YYT isolates (from YYT1) produced only one perithecium when crossed with the reference isolate, suggesting that these isolates could have retained female fertility at a very low level.

## Discussion

In this study we aimed to examinate the consequences of high levels of diversity of a cultivated plant on the associated population of a pathogen. Like many fungal pathogens, our study species is partially clonal. Sexual reproduction in *P. oryzae* populations on rice has been documented only within the worldwide lineage WW1, and is likely restricted to a few locations in South Asia. Our results confirm a previous study on *P. oryzae* in the YYT that found mating type polymorphism in the WW1 lineage while all other lineages were fixed for either mating type (Ali et al. 2023). This study provided inconsistent evidence for recombination on YYT lineages with PHI tests suggesting recombination and LD decay suggesting clonality. In our study, based on LD decay and PHI tests on an extended dataset of isolates, we found clear recombination signature only for WW1. This was expected based on previous results (Thierry et al. 2022). However, *in vitro* sexual crosses showed that all YYT isolates (including from WW1) were female-sterile and almost all were male-fertile, suggesting that even WW1 isolates have lost the ability to perform sexual reproduction and that signatures of recombination are merely inherited from the ancestral population. Although the factors determining the reproduction regime in this species are mostly unknown, it has been suggested that sexual populations of *P. oryzae* might be restricted to rainfed hillside and upland rice-growing areas (Le et al. 2023). The YYT do not fit this pattern and might be not favourable to sexual reproduction, thereby forcing clonal reproduction and, consequently, genetic loss of female then male fertility (Saleh et al. 2012). In some fungal or oomycetes pathogens, clonality is associated to invasions with, classically, low levels of diversity (e.g. Aguayo et al. 2013; Gladieux et al. 2015). In contrast our model system exemplifies a case were clonality appears to result from an environmental constraint.

The *P. oryzae* population of the YYT, despite its clonal structure, is associated with high levels of genomic diversity, mirroring the diversity of the host population. However, this diversity is mostly represented by a few but highly differentiated WW1 isolates, and a collection of unrelated clonal lineages, even if each of them harbours low levels of within-lineage diversity. The presence of such a number of distinct genetic lineages in a given location is not common in *P. oryzae.* The coexistence of the four worldwide rice blast lineages has been reported for South-East Asia (Saleh et al. 2014; Le et al. 2023) and Sub-Saharian Africa (Odjo et al. 2020; Diagne et al. 2024) and only at a regional geographical scale. Other parts of the world are generally dominated by one or two lineages (Gladieux et al. 2018b; Thierry et al. 2022). The presence of three worldwide lineages along with four apparently endemic lineages is therefore striking. Interestingly, in the mountainous region of North Vietnam (less than two hundred km distant from YYT), where numerous traditional rice landraces are cultivated, up to six different *P. oryzae* lineages were detected using microsatellite markers, four of them assigned to the four worldwide lineages and two being endemic (Le et al. 2023). Since both areas are located near the centre of origin of the species (Saleh et al. 2014), we hypothesize that large and diverse population of *P. oryzae* were introduced in these agrosystems, limiting the impact of founder effect usually observed in invasive fungal species (Gladieux et al. 2015).

We propose that incoming gene flow has only partially contributed to the current diversity of *P. oryzae* in the YYT. Ancestors of Hani people first moved into the Ailao Mountains of the Honghe River Valley in the first century AD, and adopted upland wet-rice cultivation from the 6th century (Tang Dynasty), allowing them to colonize steep mountainsides which led to the construction of the YYT along the ninth century (Cao et al. 2013; Hua & Zhou 2015). The YYT area remained fairly isolated for a long time, as the road serving the highest part was not built until the mid-20th century. This favoured the long-term maintenance of ancestral traditional indigenous practices (Jiao et al. 2012; Hannachi & Dedeurwaerdere 2018). Accordingly, our results suggested that at least some of the *P. oryzae* endemic lineages have persisted a long time in the area. Molecular clock analysis suggested that YYT1-3 have existed separately of worldwide lineages for most of the existence of the YYT. Since the YYT1-3 cluster and the YYTJ lineage were distantly related to lineages found outside the YYT, the most parsimonious hypothesis is that these two emergence events occurred in the YYT. Besides, it is interesting to note that YYTJ is potentially the first lineage to be strictly associated to japonica rice. A previous study showed that YYT isolates were organized in two groups markedly associated to indica and japonica rice (Liao et al. 2016). At the worldwide scale, WW2 isolates are preferentially associated to japonica rice (Thierry et al. 2022) but our results combined with those of Liao et al. (2016) suggest a higher level of specificity within the YYT with YYTJ lineage.

The fact that different isolates of WW2-3 (two lineages having a comparatively recent origin) were found within the YYT area, suggests that more recent immigration events also occurred. During the course of its history, the YYT underwent successive waves of settlements (Cao et al. 2013). In the 20th century, large-scale conflicts caused human population displacements, especially from Eastern China to Yunnan, and exchanges between the YYT area and the outside were facilitated by the improvement of transportation infrastructure and development of touristic activity (Gu et al. 2012; Hua & Zhou 2015). These events increased the rate of introduction of new rice varieties, including modern ones (Yang et al. 2017). As a result, the rate of introduction of non-endemic pathogen strains in the YYT has probably been increasing over time.

Although immigration of non-endemic lineages into the YYT area was likely to play a role in the current diversity of *P. oryzae*, several lineages appear to have been maintained for comparatively long time. This is somewhat unexpected, because a predominantly clonal regime of reproduction is expected to be associated with high rates of genetic drift, partly because of hitch-hiking effect caused by selection against deleterious mutations (Charlesworth et al. 1993). Furthermore, due to the elevated resistance at the population level in rice landraces in YYT (Gladieux et al. 2024), we expect that the population size of *P. oryzae* is relatively reduced. In the next two paragraph, we propose that selective factors may explain the maintenance of diversity within the pathogen population.

First, the diversity of hosts cultivated in the area maybe have triggered host specialization. In this respect, the newly described YYTJ might have had the time to specialize on japonica landraces, which are the minority in the YYT and exhibit a different repertoire of resistance genes compared to indica landraces (Gladieux et al. 2024). Like many biotrophic (including hemibiotrophic, in that case) pathogens, *P. oryzae* exhibits specialization to different host species (Gladieux et al. 2018a). However, specialization has not been observed at a lower level than rice, except in the YYT as shown already by Liao et al. (2016).

Second, selection can also maintain diversity through fluctuating selective pressures over time, e.g. in the case of negative frequency-dependent selection (Brown & Tellier 2011; Tellier et al. 2014). Farmers in the YYT switch between landraces following their own criteria (Hannachi & Dedeurwaerdere 2018; Gladieux et al. 2024) and it is likely that resistance to diseases and pests is a key factor driving their choices. Furthermore, major landraces harbour diversity, including in terms of resistance genes repertoire (De Mita et al. 2024). This system therefore paves the way toward host-pathogen coevolution and negative frequency-dependent selection (one form of balancing selection) affecting both partners.

Interestingly, the YYT1 lineage was present at a high frequency and displayed a strongly negative Tajima’s *D*. By contrast, the related YYT2 lineage exhibited an opposite pattern (low frequency and positive Tajima’s *D*). Taken together, these features are indicative of an ongoing selective sweep where YYT1 would be currently taking over YYT2. Repeated sampling over a long period of time would be needed to demonstrate the action of selection, since the changes of frequencies caused by mere genetic drift can also produce this pattern.

We conclude that the population of *P. oryzae* associated on rice landraces cultivated in the YYT has remained diversified for a prolonged period of time in spite of the relative isolation of this area during most of its history. The levels of diversity are the consequence the maintenance of different clonal lineages, including one endemic lineage that may be specialized on japonica rice. We propose that diversity has been maintained due to host specialization and/or negative frequency-dependent selection, and was incremented by human-driven introduction of worldwide lineages. This study constitutes an example of the maintenance of genetic diversity in spite of a relatively unfavourable environment caused by high levels of host resistance. From an agronomical perspective, the downside is that practices controlling the damages caused by crop diseases do not necessarily reduce the adaptive potential of the causal agent.

## Acknowledgments

We are grateful for Odile Domergue and Nicolas Poncelet for help with laboratory work, and Didier Tharreau for advices and comments. We are thankful to Pierre Gladieux, Tatiana Giraud, Romain Valade, Jean-Luc Legras and Christophe Lemaire for comments on earlier versions of the manuscript.

## Authors’ contributions

EF, HA, JBM and HH collected the samples in the YYT. EF, HA and LTL collected the samples in Vietnam. HA, LTL, SCA, SG and EF contributed to *P. oryzae* isolation and DNA extractions. PMM, SDM and SR wrote ad-hoc pipelines for genomic data curation, mapping and SNP calling. PMM, SDM and AR performed population genomics analyses. PMM, SDM and EF wrote the manuscript. All authors contributed to the revision of the manuscript. EF, HH, JBM, PMM and SDM designed the study. EF, JBM and HH provided resources for the study.

## Data availability statement

Genetic data are under process of uploading to the European Nucleotide Archive. Codes used for data analysis are available from https://gitlab.cirad.fr/phim/pmarty/rattlepm and https://gitlab.cirad.fr/phim/pmarty/scripts_vcf_to_beast

## Benefit Sharing statement

A research collaboration was developed with scientists from the countries providing genetic samples. Biological materials exchanged between collaborating laboratories were transferred under appropriate Material Transfer Agreements established between the respective institutions. These agreements defined the terms of use, ownership, and benefit sharing in accordance with institutional and applicable regulatory requirements. All collaborators are included as co-authors and the results of research have been shared with the provider communities. Benefits from this research accrue from the sharing of our data and results on public databases as described above.

## Supplementary data

**Figure S1. Population subdivision inferred with sNMF and DAPC at different K for the whole dataset (292 isolates, 41,024 SNPs).** A. Structure inferred with sNMF for K=2 to 12. Right panels show Cross-Entropy and Delta Cross-Entropy as a function of K, respectively. B. Structure inferred with DPAC for K=2 to 12. Right panels show Bayesian Information Criterion (BIC) and Delta BIC as a function of K, respectively. On bar plots, each isolate is represented by a vertical line subdivided into K segments corresponding to the probability of ancestry of the isolate in each of the K groups. Assignments of isolates established from previous studies (Gladieux et al. 2018b; Latorre et al. 2020, Thierry et al. 2022) are labelled.

**Figure S2. Population subdivision inferred with sNMF and DAPC at different K for the YYT sample (94 isolates, 20,234 SNPs).** Please refer to legend for Supplementary figure 1.

**Figure S3. Maximum-likelihood genealogy of the whole dataset (292 isolates, 41,024 SNPs) inferred with RAxML**. Isolates are coloured according their lineages attribution deduced from sNMF analyses. Dashed black lines represent YYT isolates while dashed grey lines represent isolates collected outside YYT. Scale bar represents the number of substitutions per site.

**Figure S4. Population structure inferred with genealogical network and PCA for the YYT dataset (n=94 isolates, 20,234 SNPs).** A. Genealogical network inferred with SplitsTree. Scale bar represents the number of substitutions per site. B. Principal component analysis (PCA) with the first three PC axes. Isolates are coloured according to their lineage attribution deduced from sNMF analyses.

**Figure S5. Maximum-likelihood genealogy of the YYT sample (94 isolates, 20,234 SNPs) inferred with RAxML**. Isolates are coloured according to their lineage attribution deduced from sNMF analyses. Scale bar represents the number of substitutions per site.

**Figure S6. Population tree of the YYT sample (94 isolates, 20,234 SNPs) with one migration event inferred with Treemix.** A. Population graph inferred by TreeMix (the tree is rooted with two genomes from *P. oryzae* on *Stenotaphrum*). Migration arrow is coloured according to its weight. Projections of each branch on the X-axis are proportional to the amount of genetic drift that has occurred along the branch. B. Mean and standard deviation across 10 iterations for the composite likelihood L(m) (left axis, black circles) and proportion of variance explained (right axis, red circles) as a function of the number of migration events, m. C. Second-order rate of change in likelihood (Δm) as a function of the number of migration events, m.

**Figure S7. Temporal signal in the YYT dataset restricted to non-recombining genomic regions (n=94 isolates, 17,149 SNPs).** A. Phylogenetic tree inferred with BEAST. Black circles indicate nodes at which temporal signal was found. The three nodes showing significant temporal signal and used for tip dating are pinpointed. B. Root-to-tip linear regressions for the three nodes used for tip-dating.

**Table S1. Metadata of genome accessions.**

**Table S2. Sequencing depth and coverage.**

**Table S3. Mapping depth statistics.**

**Table S4. Isolate assignation according to the literature.** Dataset compiled from Thierry et al. (2022), Ali et al. (2023) and sNMF and DAPC results for this study.

**Table S5. Statistics per pair of populations.** F_ST_WC: Weir and Cockerham’s F_ST_. Dj: Jost’s differentiation index. D_a_: net pairwise distance. D_xy_: absolute pairwise distance. S: number of variant sites. lseff: number of sites analysed.

**Table S6. Results of PHI test per lineage.** Only lineages with at least 3 isolates were considered.

