## Supplementary figures and images for "Population structure of the rice blast fungus *Pyricularia oryzae* in the context of elevated host heterogeneity"

### Supplementary Figure 1

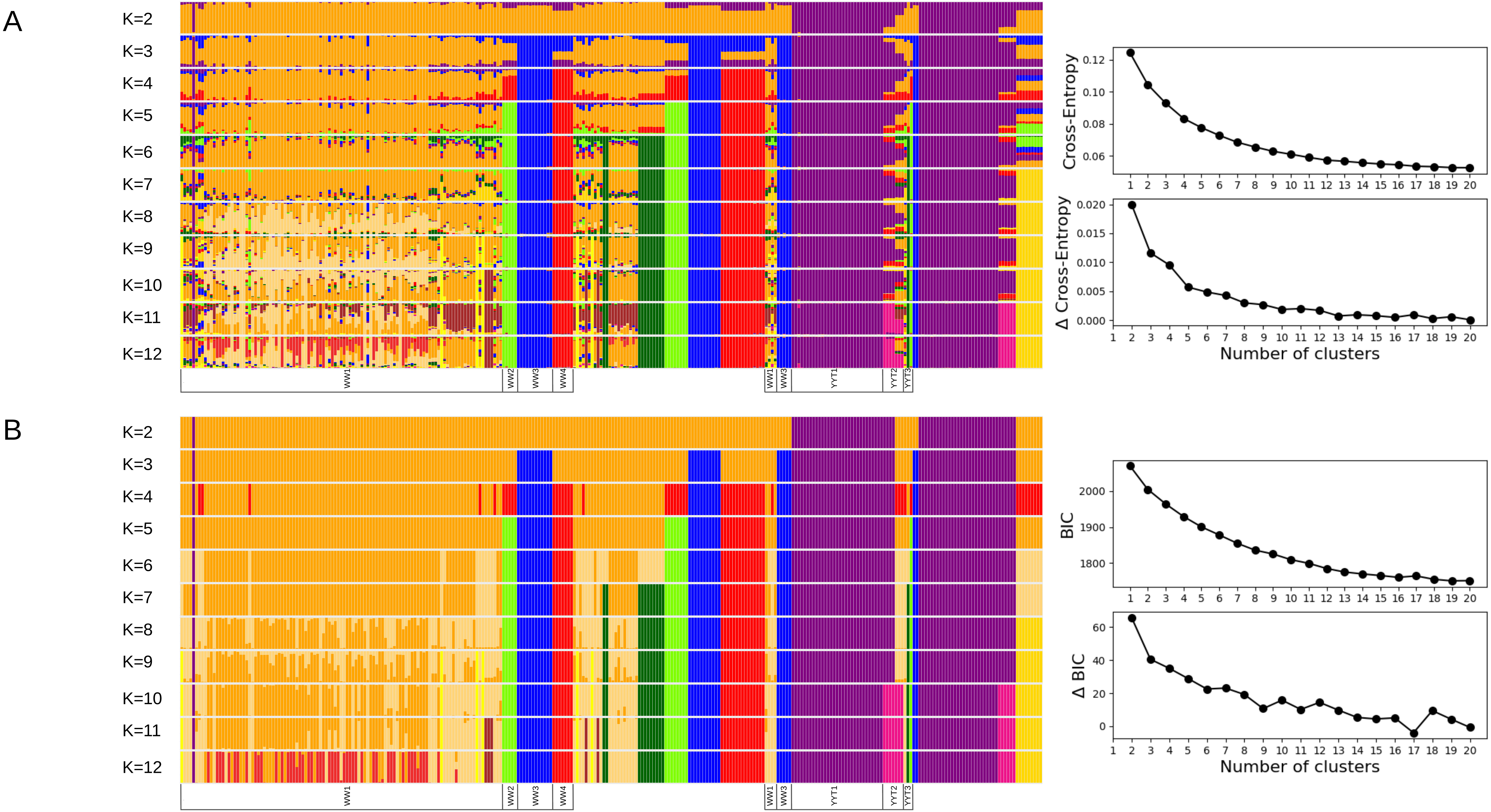

### Supplementary Figure 2

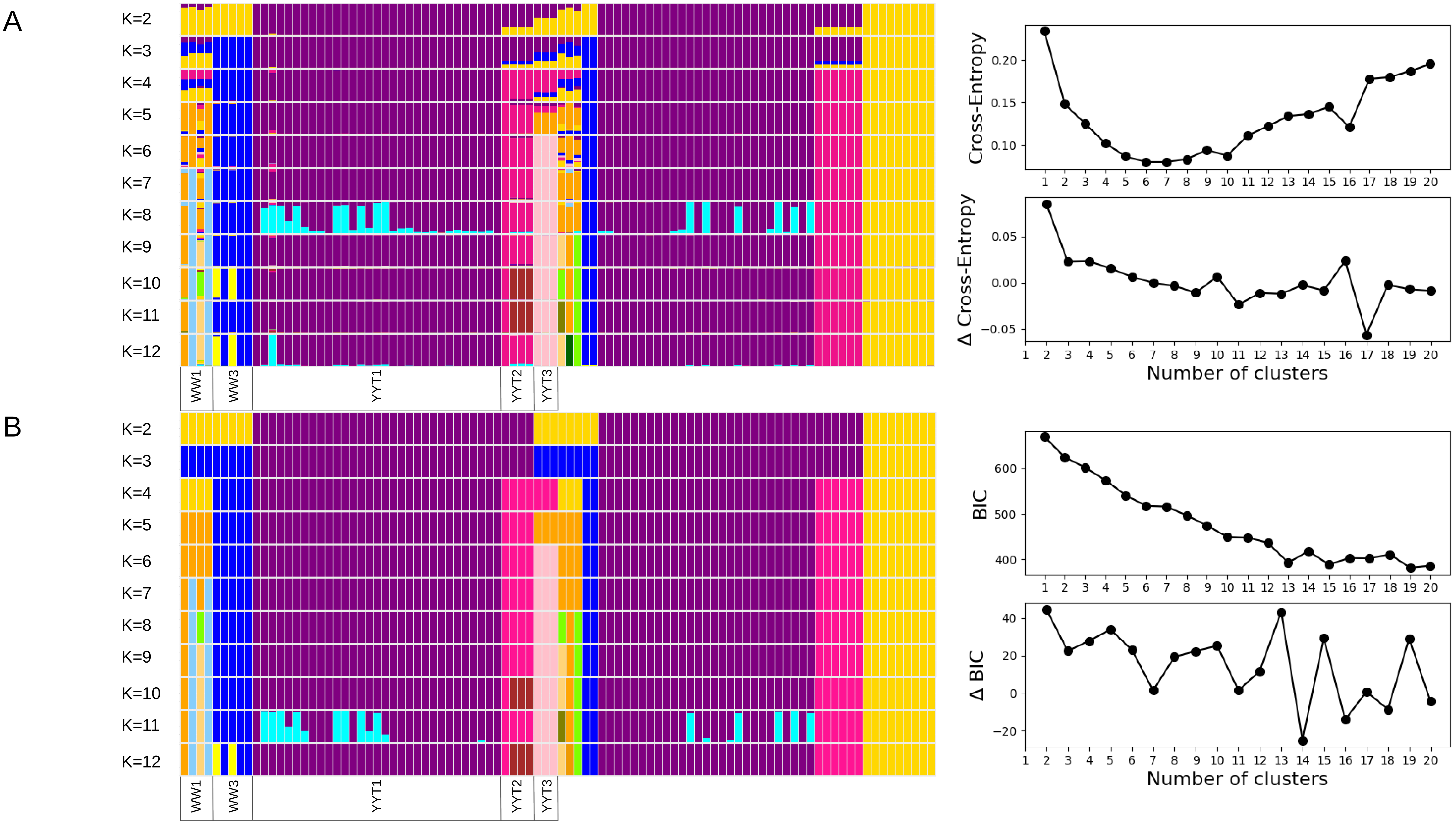

### Supplementary Figure 3

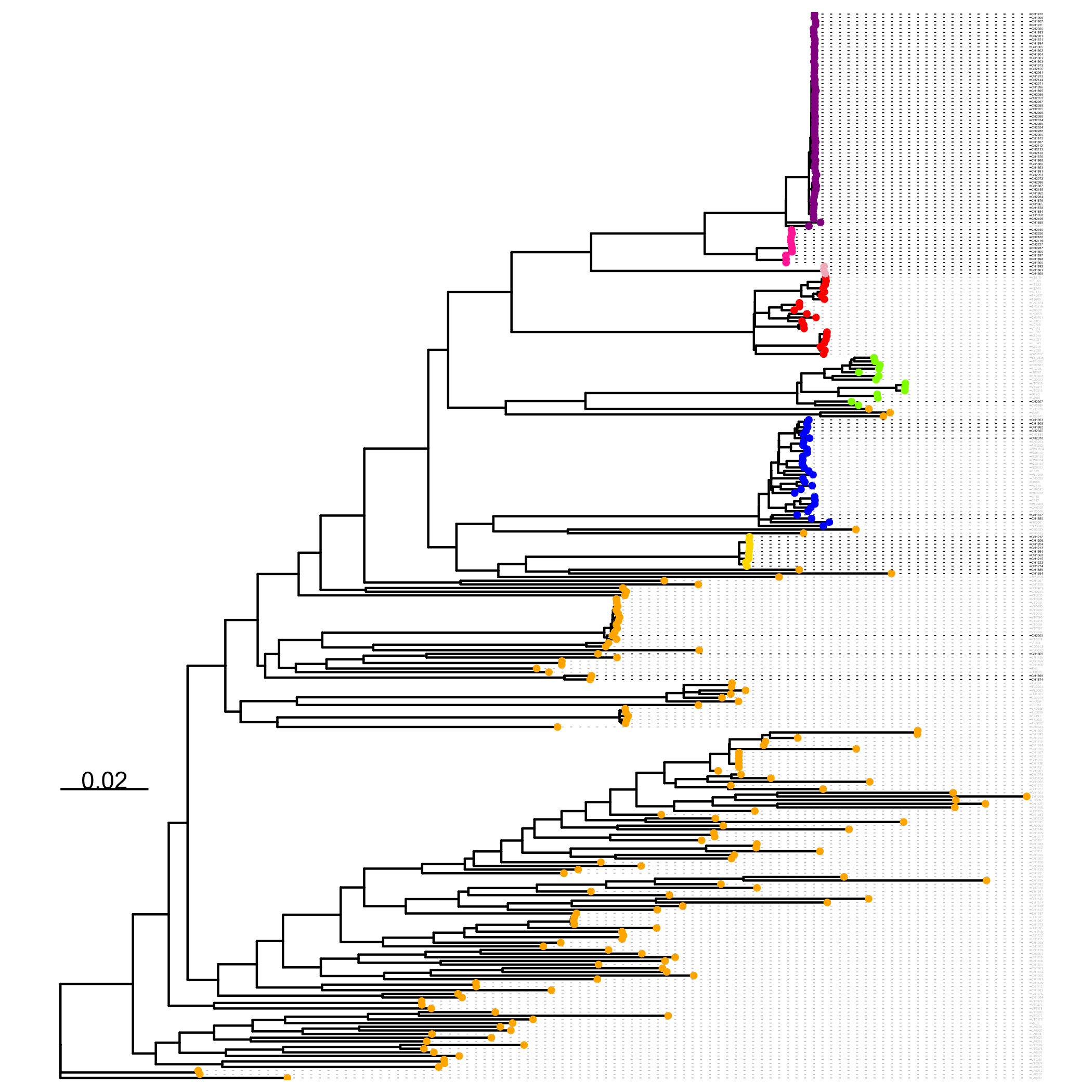

### Supplementary Figure 4

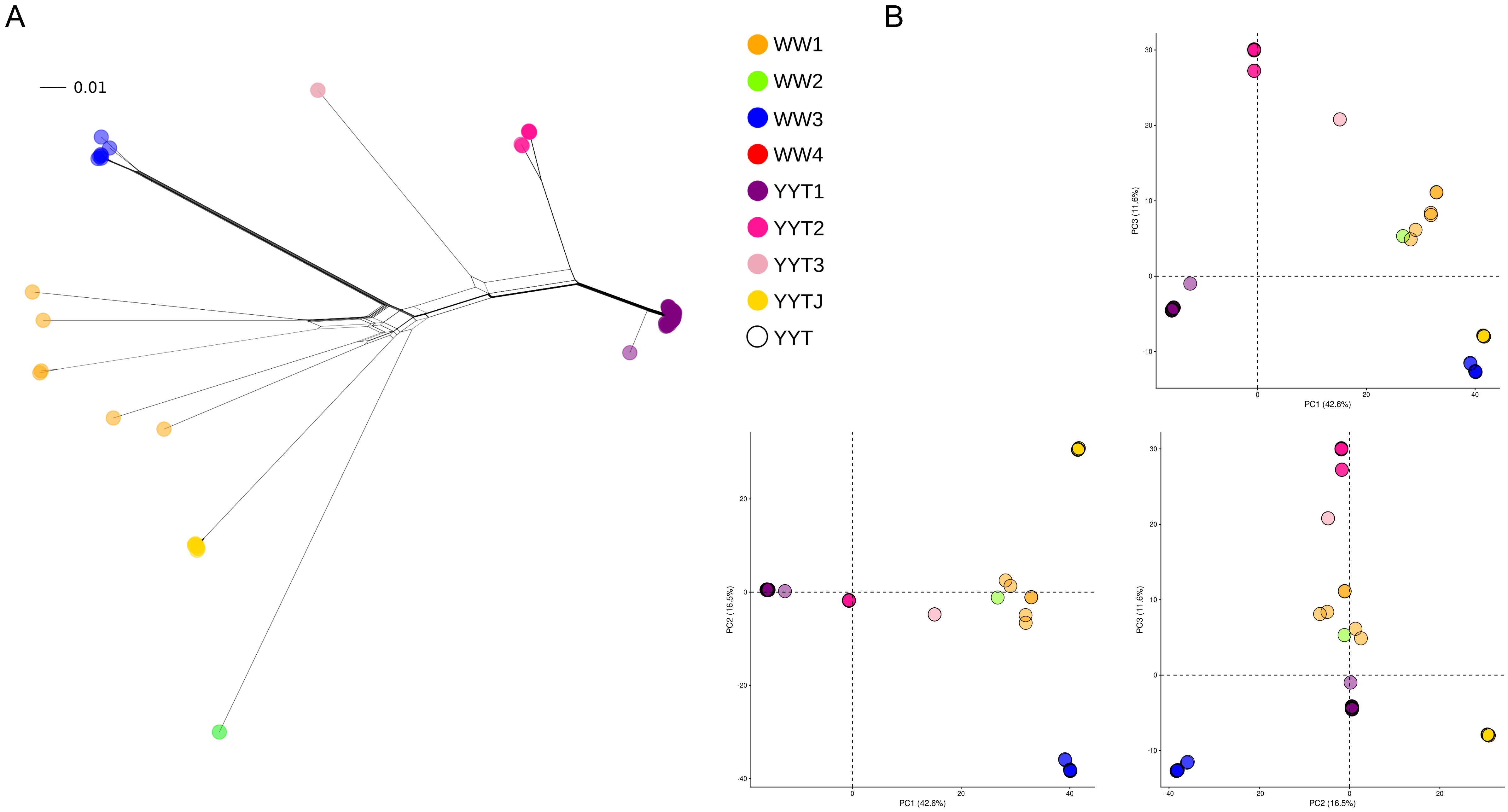

### Supplementary Figure 5

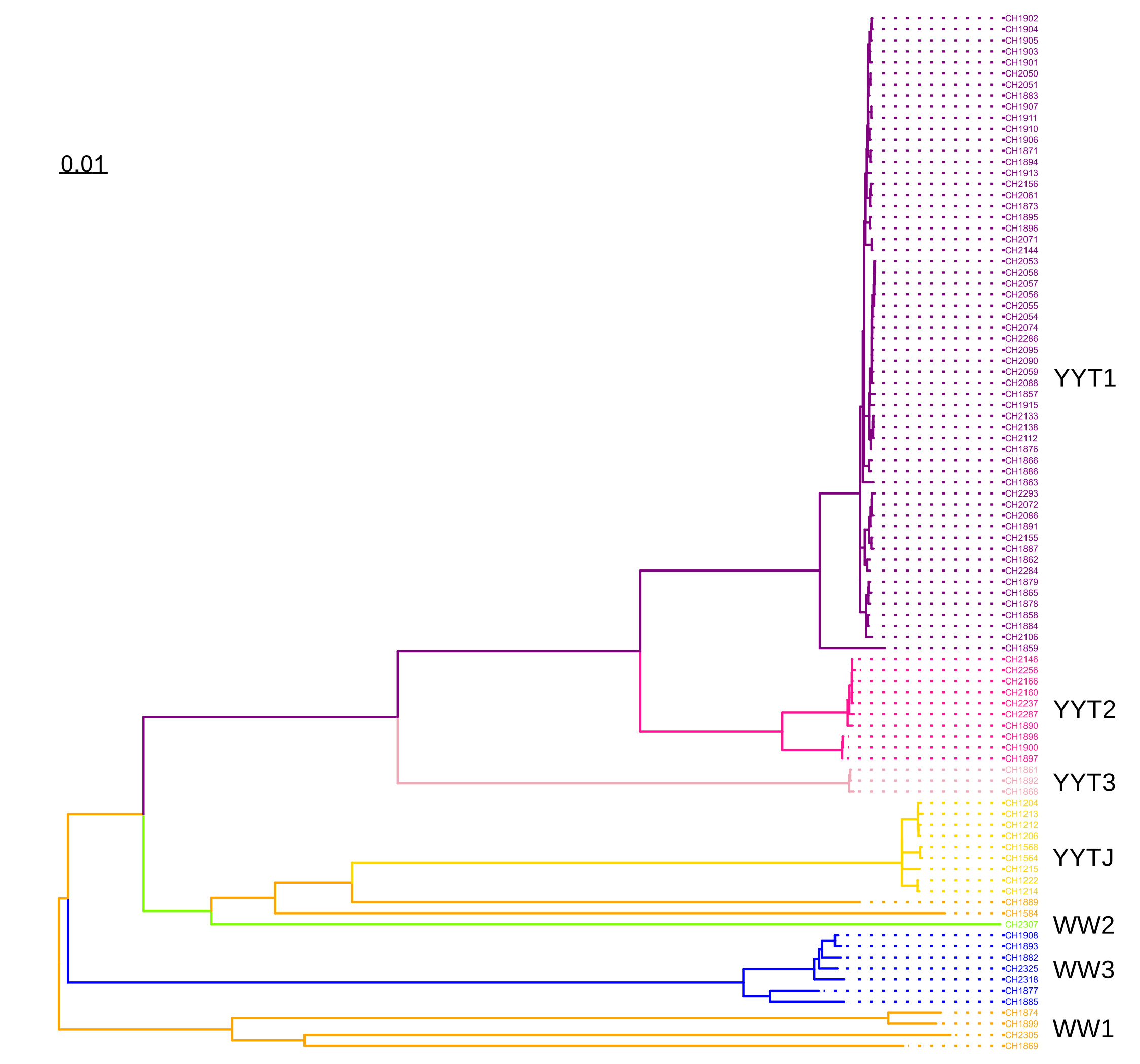

### Supplementary Figure 6

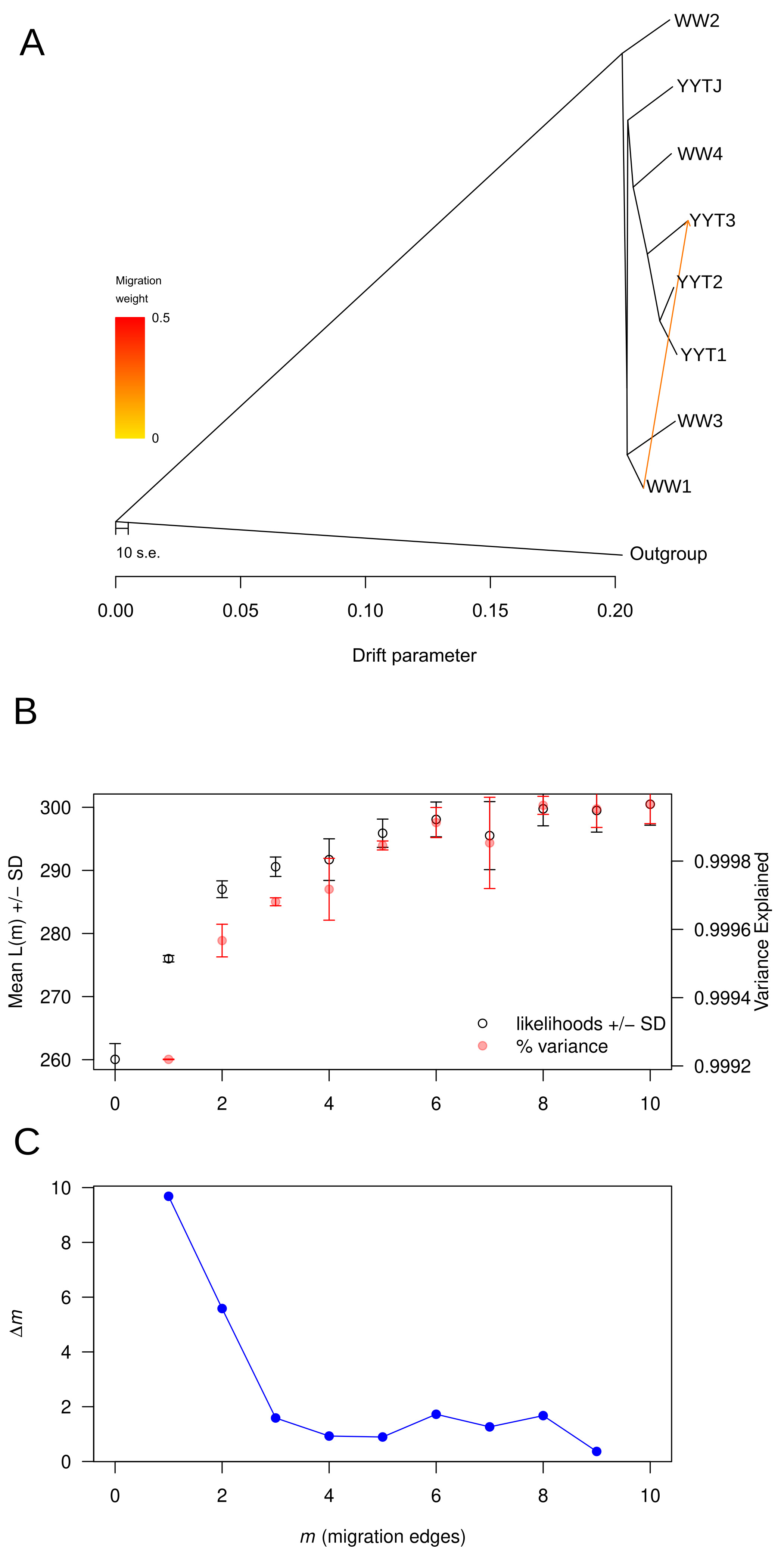

### Supplementary Figure 7

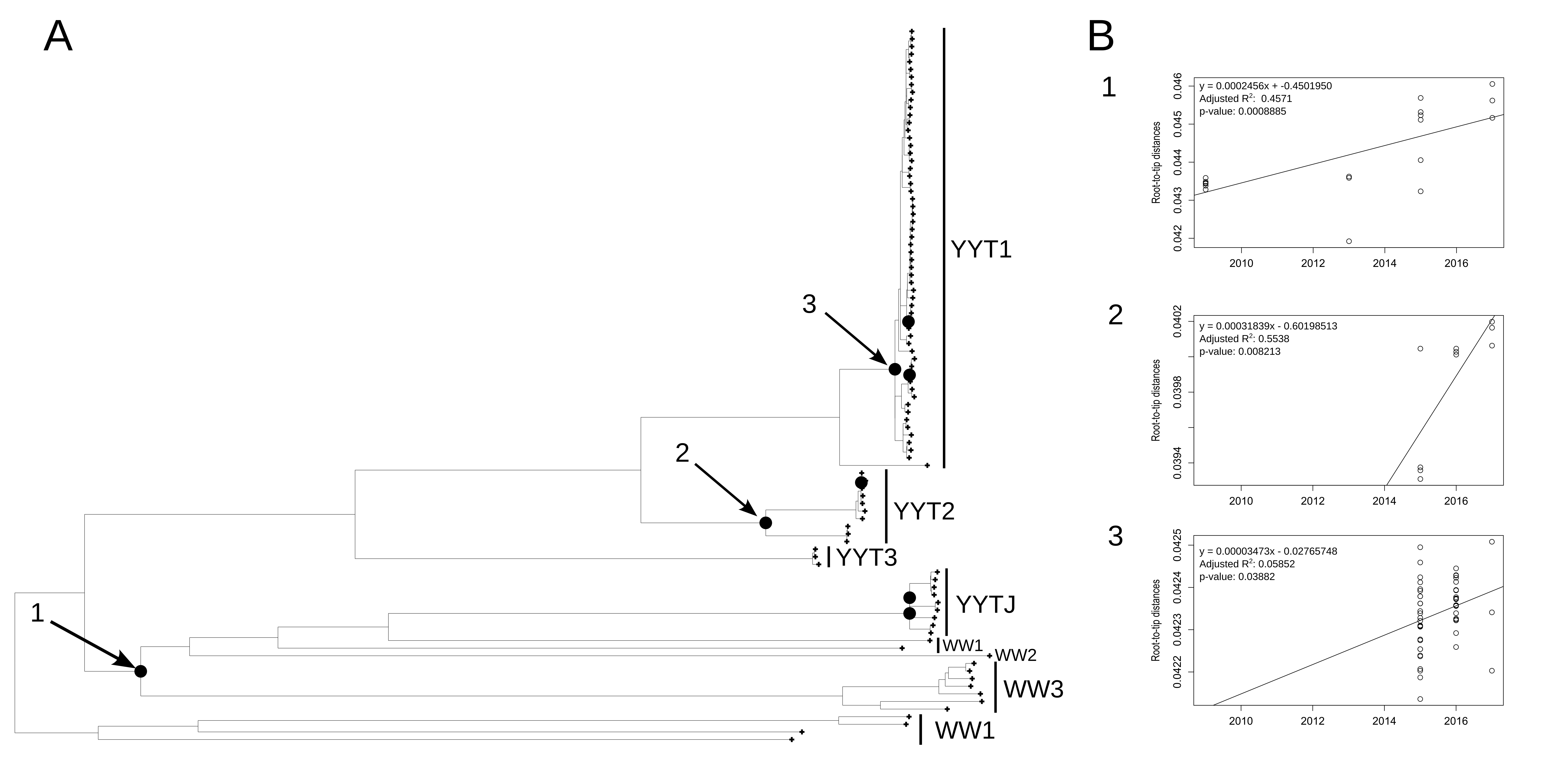
